# Acod1 negatively impacts osteoclastogenesis via GPR91-mediated NFATc1 activation

**DOI:** 10.1101/2022.04.07.487467

**Authors:** Yue Gao, Franziska V. Kraus, Elisabeth Seebach, Sushmita Chakraborty, Dominik Schaack, Judith Schenz, Willi Wagner, Katharina F. Kubatzky

## Abstract

Immune cells undergo metabolic reprogramming upon exposure to inflammatory stimuli. The immuneresponsive gene 1 (*Irg1*) encodes aconitate decarboxylase (Acod1), an enzyme that generates itaconate from cis-aconitate in the TCA cycle and is one of the most highly induced genes in macrophages during inflammation. Itaconate inhibits succinate dehydrogenase activity leading to the accumulation of succinate. As the adjustment of energy metabolism also plays an important role during the formation of bone-resorbing osteoclasts, we investigated if *Irg1* is regulated during osteoclastogenesis. We show that M-CSF/RANKL treatment induces *Irg1* at an early time-point in bone marrow-derived macrophages (BMDM) as well as in the RAW264.7 macrophage cell line. Next, we stably overexpressed Acod1 in RAW264.7 cells. The metabolism of these Acod1 cells shifted towards glycolysis, as indicated by an increase in mTOR activation, subsequent 4EB-P1 phosphorylation and reduced ATP levels. When we investigated the ability of Acod1 cells to differentiate into osteoclasts, we observed a remarkable suppression of osteoclast-associated genes and the number of TRAP-positive, multi-nucleated osteoclasts was greatly reduced but not completely abrogated. Surprisingly, NFATc1 was detectable in nuclear extracts in untreated Acod1 cells leading to residual transcriptional NFAT activity in luciferase assays. This is caused by the elevated levels of succinate in Acod1 cells, as succinate can bind extracellularly to its cognate receptor GPR91 leading to Gq-mediated activation of NFATc1. When we investigated the expression of *Gpr91*, we found RANKL-mediated induction of *Gpr91* to be severely reduced in Acod1 cells and we suggest that GPR91 is a target of RANKL-mediated NFATc1 activation. However, on the protein level, the receptor was still expressed at the cell surface. The observed repression of *Gpr91* in Acod1 overexpressing cells was also detected by treatment with octyl-itaconate, showing that this is an itaconate-mediated effect. We hypothesize that the itaconate-mediated increase in succinate levels causes activation of NFATc1 signalling, although the transcriptional activity does not lead to osteoclastogenesis. In the presence of RANKL, these pre-activated cells are slow in switching to RANKL-mediated induction of osteoclast genes, which decreases their ability to differentiate into osteoclasts.

## Introduction

Osteoclasts are derived from haematopoietic cells and have the unique ability to resorb bone. As immune cells, they can also interact with T cells and shape the immune response by acting as antigen presenting cells (1). Under *in vitro* conditions, the differentiation of osteoclasts is driven by the two cytokines macrophage colony stimulating factor (M-CSF) and receptor of activated NF-kB ligand (RANKL). However, other factors such as hormones, pro-inflammatory cytokines or bacteria-derived substances can contribute directly or indirectly to osteoclast formation and function (2). Especially under inflammatory conditions, macrophage activity and osteoclast differentiation mutually influence each other (3). Cytokines, such as IL-6, TNF-α and IL-1β, are primarily pro-inflammatory stimuli, but also have a profound effect on osteoclastogenesis leading to the disbalance in bone homeostasis observed in many chronic inflammatory diseases (4). In recent years, the influence of the metabolic regulation of macrophage function, activity and plasticity came into focus, however, the impact of metabolism on osteoclast differentiation is less well studied. Due to the connection between inflammation and osteoclast activation, a better biochemical understanding of metabolic aspects of the macrophages and osteoclasts are needed.

Quiescent cells preferentially produce ATP via the TCA cycle and oxidative phosphorylation (OxPhos) to guarantee maximal production of energy from glucose, however, activated immune cells are able to adapt their metabolism to the required function (5, 6). Although glycolysis generates a lower amount of ATP, it creates more building blocks that can then quickly go into anabolic pathways. For macrophages, it was shown that the change into proinflammatory M1-type macrophages triggered by TLR ligands and IFN-γ is coupled to a metabolic switch towards glycolysis, while anti-inflammatory M2-type cells prefer OxPhos. Bone resorption is an energy demanding process and thus the metabolic activity is a central aspect in the osteoclast differentiation process, where both glycolysis and TCA cycle progression are needed (7). Jin et al. demonstrated that the Ndufs4 protein, a subunit of the complex I of the ETC, is a central switch between macrophage activation and osteoclast differentiation (8). In the absence of Ndufs4, ETC activity and the TCA cycle are disrupted, so that the cell is forced to switch towards glycolysis. Because osteoclast differentiation is not possible in the absence of a functional ETC, this enforced metabolic shift results in systemic inflammation and an osteopetrotic phenotype with increased bone density. This suggests that the metabolic activity initiates a change in cellular function and not vice versa. Immune reactions can also change the output of metabolites. Metabolites that accumulate during glycolysis and the TCA cycle play a role in macrophage activation and osteoclastogenesis (9, 10). Cis-aconitate is a by-product when citrate is converted into iso-citrate and its production can result in anti-inflammatory signalling as aconitate is the target of aconitate decarboxylase (Acod1), the enzyme that produces itaconate. Acod1 was first discovered as an “immune responsive gene” 1 (*Irg1*) due to its massive upregulation after LPS stimulation (11) but it took nearly twenty years to discover its enzymatic function and its importance as a regulator of the TCA cycle and the inflammatory state of macrophages (12). Because the absence of Acod1 prolongs cytokine production, it was suggested that itaconate has an anti-inflammatory function (13). In addition to diverting citrate out of the TCA cycle, itaconate has the ability to block succinate dehydrogenase (SDH), so that succinate accumulates. Metabolically, Acod1-deficient cells therefore display enhanced OxPhos activity, fatty acid oxidation and glutaminolysis (14, 15). Acod1 overexpression on the other hand limits the LPS-mediated production of cytokines such as TNF-α, IL-6 and IL-1β (16). Succinate on the other hand seems to be a pro-inflammatory mediator in inflammatory settings. When succinate is processed by SDH, the resulting ROS helps to stabilise HIF-1α eventually resulting in the production of IL-1β (17). Itaconate-mediated inhibition of SDH during macrophage activation causes accumulation of succinate in the cytosol where it can change protein activation and transcription factor activity through succinylation of proteins, histones or DNA (18). When succinate accumulates, it can also be excreted to the extracellular space where it binds to the succinate receptor (SUNCR1/GPR91) (19). GPR91 is a Gαi/Gαq-dependent GPCR that is expressed on haematopoietic cells and plays a role in pathologies like obesity, diabetes, hypertension or rheumatoid arthritis (RA) where increased levels of extracellular succinate can be observed. Succinate is discussed to be a valuable biomarker and a potential target for disease therapy (20). The effect of succinate-induced signalling in the absence of an already primed, (pro- or anti-inflammatory) microenvironment is currently unclear. GPR91 is also expressed on cells of the osteoclast lineage and extracellular succinate was found to enhance osteoclastogenesis (21, 22). As a consequence, diabetes patients have an underlying bone impairment because of increased osteoblast apoptosis and overactive osteoclasts. Correspondingly, the addition of RANKL/M-CSF together with succinate increased the number and activity of osteoclasts in an *in vitro* assay. This was also observed *in vivo*, where mice that had been injected with succinate showed significant bone loss. Metformin, a drug frequently used to treat type 2 diabetes, is a complex I inhibitor that resets the metabolite balance, reduces succinate levels and ultimately blocks excessive osteoclast formation (22). These data emphasize that especially under pathological conditions a vicious circle can occur that connects macrophage reprogramming and osteoclast activity. In auto-inflammatory diseases, a disease-specific shift in metabolic activity can be observed (23). As succinate accumulation in the synovial fluid increases the inflammatory phenotype of macrophages in RA (22) and hypermetabolic macrophages were discovered in RA and coronary artery disease (24), it can be hypothesized that Acod1 and metabolic changes in macrophages in general, might have an impact on inflammatory and auto-immune diseases. Here, we show that Acod1 activity suppresses osteoclast formation. The enhanced levels of succinate in Acod1 overexpressing cells caused GPR91 activation and subsequent signalling through the Gq subunit of the associated heterotrimeric G protein. In the lack of a permissive microenvironment, the resulting NFATc1 activity was non-productive for the induction of osteoclast genes and seemed to decrease the sensitivity of the cells for RANKL-mediated signal transduction. In addition, itaconate is a regulator of *Gpr91* gene expression, but the more important step of succinate regulation seems to occur at the protein level. We found that the availability of GPR91 for succinate signalling is decreased by Acod1 overexpression as well as RANKL treatment, presumably to limit osteoclastogenesis.

## 2. Materials and Methods

### 2.1 Mice

C57BL/6 wild-type mice were purchased from Janvier Labs (LeGenest St. Isle, France), and Acod1-deficient Acod1^em1(IMPC)J^/J mice (JAX stock #029340) were obtained from Jackson Laboratories (Bar Harbor, ME, USA). Mice were maintained under SPF conditions and killed in accordance with the German policies on animal welfare.

### 2.2 Reagents

Tissue culture reagents were purchased from Anprotec (Bruckberg, Germany), Biochrom GmbH (Berlin, Germany), PAN biotech (Aidenbach, Germany), Thermo Scientific (Langenselbold, Germany), Merck and Sigma. Antibodies against phosphorylated NF-kB (Ser536), AMPKα (Thr172), 4E-BP1 (Thr37/46), total IRG1, HSP90, Lamin B1, BLIMP1, and NFATc2 were purchased from Cell Signaling Technology (Leiden, Netherlands). Total NFATc1 was purchased from BD Pharmingen (San Diego, CA, USA). Antibodies against GAPDH and β-actin were obtained from Proteintech (Manchester, UK). Secondary horseradish peroxidase (HRP)-linked antibodies were obtained from Cell Signaling Technology. PCR primers were purchased from Biomers (Ulm, Germany).

### 2.3 Lentiviral transduction

To generate an overexpressing Acod1 cell line, a total of 300,000 of RAW 264.7 (ATCC^®^ TIB-71™) were seeded in 2 mL of DMEM medium containing 10% FCS and 1% Pen/Strep in 6-well plates. Cells were incubated at 37 °C and 5% CO_2_ overnight before transduction. Recombinant lentiviral vectors (Acod1: pLV[Exp]-EGFP:T2A:Puro-EF1A>HA/mAcod1[nm_0008392.1] Mock: pLV[Exp]-EGFP:T2A:Puro-EF1A>mCherry) purchased from VectorBuilder (Chicago, IL, USA) were added at a MOI of 10 into the cells followed by 1 h spin infection at a speed of 300 g. Puromycin selection was performed after 48 h incubation in fresh medium. Transduction efficiency was verified by flow cytometry.

### 2.4 Differentiation and stimulation of cells

To generate primary murine macrophages, bone marrow (BM) cells were isolated from the femur, tibia and humerus of the over 10 weeks-old C57BL/6 mice. Cells were treated with L929-cell conditioned medium (LCCM) as described previously (25). On day 1, BM cells were resuspended in 20 ml of DMEM medium (with 10% FCS, 1% Pen/Strep and 0.1% 2-Mercaptoethanol). On day 4, cells were restimulated with 30% LCCM and incubated for additional 3 days before seeding and performing experiments. Cells were cultivated in DMEM medium and stimulated with 10 ng/mL LPS from *Salmonella enterica* diluted from 1 mg/mL stock (L5886 Sigma-Aldrich, Taufkirchen, Germany) with culture medium. 100 mM 4-octyl Itaconate (Cayman Chemical Company, Ann Arbor, US) from the DMSO dissolved stock (AppliChem, Darmstadt, Germany) were added to the cells 3 h prior to the desired stimulation. For osteoclast differentiation, cells were stimulated with 50 ng/ml rec. mouse sRANKL, BMDMs were supplemented with additional 25 ng/ml rec. mouse M-CSF (all from Biotechne, Abington, UK).

### 2.5 Quantitative real-time PCR

A total of 80,000 Raw cells (numbers were decreased into half as days proceed) or 200,000 BMDMs were seeded and stimulated on a 24-well plate as indicated. RNA-isolation was carried out either by innuPREP RNA Mini Kit 2.0 from Analytik Jena (Jena, Germany) or Extractme Total RNA Micro Spin Kit from 7Bioscience (Neuenburg, Germany) according to the manufacturer’s protocol, respectively. cDNA synthesis was performed subsequently by applying Biozym cDNA Synthesis Kit (Biozym Scientific GmbH, Hessisch Oldendorf, Germany). Quantitative real-time PCR was performed by using qPCRBIO Syber Green Mix Hi-ROX (PCR Biosystems, Wayne, US) and run with the StepOne Real-Time PCR System (Applied Biosystems, Waltham, US). The relative gene expression was calculated as 1/2^ΔCT^ based on the normalization by the reference gene *Hprt1*. For human samples, RNA was extracted using GeneJET RNA Purification Kit (Thermo Fisher Scientific, USA), according to the manufacturers’ protocol. cDNA was prepared by using Revert Aid First strand cDNA synthesis kit (Thermo Scientific). Quantitative RT-PCR was performed using Powerup Sybr Master Mix (Applied Biosystem) with the primers mentioned below. RT-PCR was performed using the QuantStudio 5 Real-Time PCR Systems (Applied Biosystems). An initial enzyme activation step of 2 min at 50°C and denaturation step of 5 min at 95°C, followed by amplification for 40 cycles at 95°C for 15 s and at 58°C for 1 min. As normalization control *GAPDH*, was used. All primers used are stated in the supplementary table.

### 2.6 Cell separation and cell culture

Synovial Fluid mononuclear cells (SFMC) and Peripheral blood mononuclear cells (PBMCs) were isolated using Lymphoprep (Axis-Shield, Oslo, Norway) density gradient centrifugation. Cells were resuspended in RPMI-1640 containing L-Glutamine, and HEPES (Gibco) supplemented with, 10% FBS (Gibco) and 100 U/ml penicillin, 100 μg/ml streptomycin (Sigma-Aldrich, St. Louis, MO, USA). A total of 5 × 10^6^ cells were seeded per well of 6 well plate. Cells were kept in humidified 5% CO_2_ incubator at 37 °C for 24 hours. After 24 hours, non-adherent cells were removed by washing the wells with PBS three times. Adherent cells were processed for expression studies.

### 2.7 Mitochondrial copy number

A total of 100,000 RAW264.7 cells were seeded and stimulated on a 24-well plate as indicated. On day 4, the cells were harvested and the total genomic DNA was isolated by using the innuPREP DNA mini kit (Analytik Jena, Jena, Germany). 10 ng of DNA was used for amplification and quantification according to a qPCR approach. The mitochondria copy number was evaluated by comparing the ratio between mitochondrial DNA amount and nuclear DNA amount (mtDNA/nucDNA) in the cells.

### 2.8 Cell Viability Assay

A total of 50,000 cells were seeded as triplicates in a translucent 96 well microplate and stimulated as indicated. To quantify ATP levels, CellTiter Glo^®^ reagent (Promega, Mannheim, Germany) was added on the well after removing supernatants. For the measurement, cells lysates were transferred to a white 96 well assay plates and evaluated by Luminometer LUMIstar Optima (BMG, Offenburg, Germany).

### 2.9 FACS Analysis

For detection of extracellular markers, cells were blocked in PBS, 2% BSA for 15 min on ice in a total volume of 100 μL. Cells were then incubated with the primary antibody (Anti-GPR91, Alomone labs, Jerusalem, Israel) for 1 h. The staining step was carried out with allophycocyanin cross-linked antibody (BioLegend, San Diego, US). For RANK expression, cells were incubated with the allophycocyanin cross-linked anti-RANK antibody (BioLegend, San Diego, US). Surface expression was quantified by flow cytometry by using FACS Canto cytometer (BD Biosciences, Heidelberg, Germany) and BD FACS Diva Software. For the live/dead staining, cells were stained with Zombie NIR™ Fixable dye (BioLegend, San Diego, US) diluted as 1:10 in PBS from DMSO stock for 15 min. Cells were analysed within the APC channel after PBS washing. For measuring mitochondrial activity and oxidative stress, the cells were stained with 100 nM MitoTracker™ Deep Red (Cell Signaling Technology, Leiden, The Netherlands) and 5 μM CellROX™ Deep Red (Thermo Fischer Scientific, Waltham, US), respectively. After 30 min incubation, the cells were washed with PBS and analysed within the APC channel.

### 2.10 Western Blot Analysis

For Raw264.7 cells, a total of 800,000 cells (numbers were decreased into half as days proceed) were seeded and stimulated on a 12-well plate as indicated. For BMDMs, a total of 2,000,000 cells were seeded and stimulated on 6-well plate as indicated. After PBS washing, cells were lysed with 200 μl of 1 × RIPA buffer, freshly supplemented with PhosSTOP™ inhibitor tablets for phosphatase and cOmplete™ Protease-Inhibitor Cocktail (Roche, Basel, Switzerland). 10 μg of protein per sample were collected and normalized according to BCA assays. Samples were run on an SDS-PAGE 4–20% gradient polyacrylamide gel (Anamed, Gross-Bieberau, Germany). Proteins were transferred to nitrocellulose membranes (GE, Boston. US) via 1 h semi-dry blot. Blotted membranes were blocked with 1x Blue Block PF blocking buffer (SERVA) for 30 min at RT, followed by overnight primary antibody incubation at 4°C. After 1 h incubation with HRP coupled secondary antibody, protein bands were detected by enhanced chemiluminescence (Intas Science Imaging, Göttingen, Germany).

### 2.11 TRAP Assay

To evaluate osteoclast differentiation, a total of 5,000 RAW cells were seeded and stimulated on a 24-well plate. For BMDMs, a total of 70,000 cells were seeded and stimulated on a 48-well plate. Restimulation was applied on day 3 by replacing half volume of medium with fresh medium and stimuli. Cells were fixed and stained for approximately 30 min by applying Acid Phosphatase, Leukocyte (TRAP) Kit (Sigma-Aldrich, Burlington, US) at 37°C. TRAP-positive cells with three or more nuclei were counted as osteoclasts.

### 2.12 Gene reporter Luciferase Assay

A total of 200,000 RAW cells were seeded on a 24-well plate and were co-transfected with NFATc1(pGL3-NFAT) and *Renilla* luciferase plasmid (pRL-TK, Promega) in a ratio of (2:1), supplemented with 2 μl of Lipofectamine™ 2000 (Invitrogen, Waltham, US) per sample (26). After 4 h Cells were stimulated with 500 μM sodium succinate (Sigma-Aldrich, Taufkirchen, Germany) for 6 h. The transcriptional activation was evaluated by using the Dual-luciferase^®^ reporter assay system (Promega, Mannheim, Germany). Cell lysates were then transferred to a white 96-well assay plate and evaluated by Luminometer LUMIstar Optima (BMG, Offenburg, Germany).

### 2.13 μCT scans

To analyse possible changes in bone density or size in the knock out (k/o) mice compared to the wild type (WT) mice, μCT scans were performed using a Sky-Scan 1176 in vivo X-ray microtomograph (Bruker Corporation, USA). As the bone properties differ in males and females and are also changing in an age dependent manner, analysed bones were used from age- and sex-matched mice (all males, ≥ 10 weeks old). Upon dissection, femora and tibiae were fixed in 4% paraformaldehyde for at least 24 h. In order to prevent minimal movement of the samples during the scanning, the bones were embedded in 1% agarose in water one day prior the measurement. The samples were placed in axial direction for the scan using the 0.5 mm aluminium filter, with the following settings: voxel size 17.3 μm, voltage 50 kV, current 500 μA, exposure time 225 ms, frame averaging 4. Data were recorded every 0.5° rotation step through 180°. Reconstruction was performed using Skyscan NRecon^®^ software (version 1.7.3.0, Bruker Corporation, USA). 3-D pictures were made with Skyscan CTan^®^/CTVol^®^ software (Bruker Corporation, USA). Bone parameters were analysed with Skyscan CTan^®^ analysis software (Bruker Corporation, USA). To analyse comparable volume of interest (VOI) in all samples, landmarks were chosen: in the femora samples, 150 images in distal direction from the trochanter tertius were selected for analysis. For the tibiae samples the first image was selected, where no connection to the fibula could be observed anymore. From there on, 150 images in proximal direction were included in the analysis. For evaluating bone density, the mean greyscale index within the VOI was determined. For analysis of bone volume per tissue volume (BV/TV) and bone surface per bone volume (BS/BV) the lower grey level was set at 70 and the upper grey level was set at 255. μCT-analysis was performed according the guidelines (27) and results are reported following ASBMR nomenclature (28, 29).

### 12.14 Study Subjects

The study was conducted with approval from Institute Ethics Committee **(IEC-490/01.09.2017)**. After obtaining informed consents, 8 patients with RA and OA, each, were recruited from Orthopaedics OPD of AIIMS, New Delhi, India. All recruited RA patients had active disease. For this study, we collected synovial fluid (SF) and autologous peripheral blood (PB) from RA patients. Peripheral blood from OA was collected. Specimens were collected in heparinized tubes (BD, USA).

### 2.15 Data processing and statistical analysis

The BD FACS Diva software and Flowing Software 2 were used for the analysis of flow cytometry data. Statistical analysis and graph design were performed using Graph Pad Prism version 8 (San Diego, CA, USA). Comparison between 2 groups was performed as indicated in the figure legend for each experiment.

## 3. Results

### 3.1 RANKL induces Acod1 expression

LPS and other TLR ligands are well-known inducers of *Irg1*. In addition, various pro-inflammatory cytokines, such as IFN-β and TNF-α are able to induce Acod1 expression (30, 31). As it was shown previously that octyl-itaconate can inhibit osteoclastogenesis (32), we investigated, whether the cytokine RANKL which is a member of the TNF family, is able to induce the production of endogenous itaconate through *Irg1* expression. We therefore stimulated primary bone marrow-derived macrophages as well as RAW264.7 macrophages with M-CSF, M-CSF/RANKL or LPS as a control and determined the gene transcription of *Irg1*. As an initial time-course between 3 and 24 hours indicated a strong induction at 6 hours (Figure S1A), we focussed at the 6 h time-point in our further experiments (Figure 1A and B). In both cell types we observed a significant increase in *Irg1* levels, however, the effect was more pronounced in the cell line. Next, we created Acod1 overexpressing RAW264.7 cells (Acod1 cells) by transducing them with the lentiviral vector pLV[Exp]-EGFP:T2A:Puro-EF1A>HA/mAcod1 or the empty vector as a control (Mock cells). After puromycin selection, the transduction efficiency was determined by measuring eGFP expression which corresponds to the level of Acod1 (Figure 1C). The result was confirmed by checking the gene expression of *Irg1* in untreated cells. Figure 1D shows a significant overexpression of the gene in untreated cells. To confirm that the overexpressed Acod1 is functional and creates the expected phenotype, we investigated ATP production after 4 and 24 h (Figure 1E). As Acod1-produced itaconate blocks the TCA cycle, overexpression of Acod1 should result in lower ATP levels. Figure 1E demonstrates that ATP levels are indeed decreased at both time-points. To ensure that this is neither caused by nor leads to increased cell death, we performed a life-dead stain. Figure 1F shows that the percentage of viable cells does not differ significantly between Mock and Acod1 transfected cells. In addition, we observed an upregulation of the mTOR pathways as a marker for glycolytic activity, which was accompanied by an increase in phospho adenosine monophosphate kinase (AMPK) (Figure S1B). AMPK is a sensor of low ATP levels, that when activated, helps the cells to switch to TCA activity and OxPhos-mediated ATP production. This suggests that due to the enhanced glycolytic activity and the decrease in ATP production observed in Figure 1E, the cells attempt to switch back to TCA cycle activity by AMPK phosphorylation.

**Figure 1:**
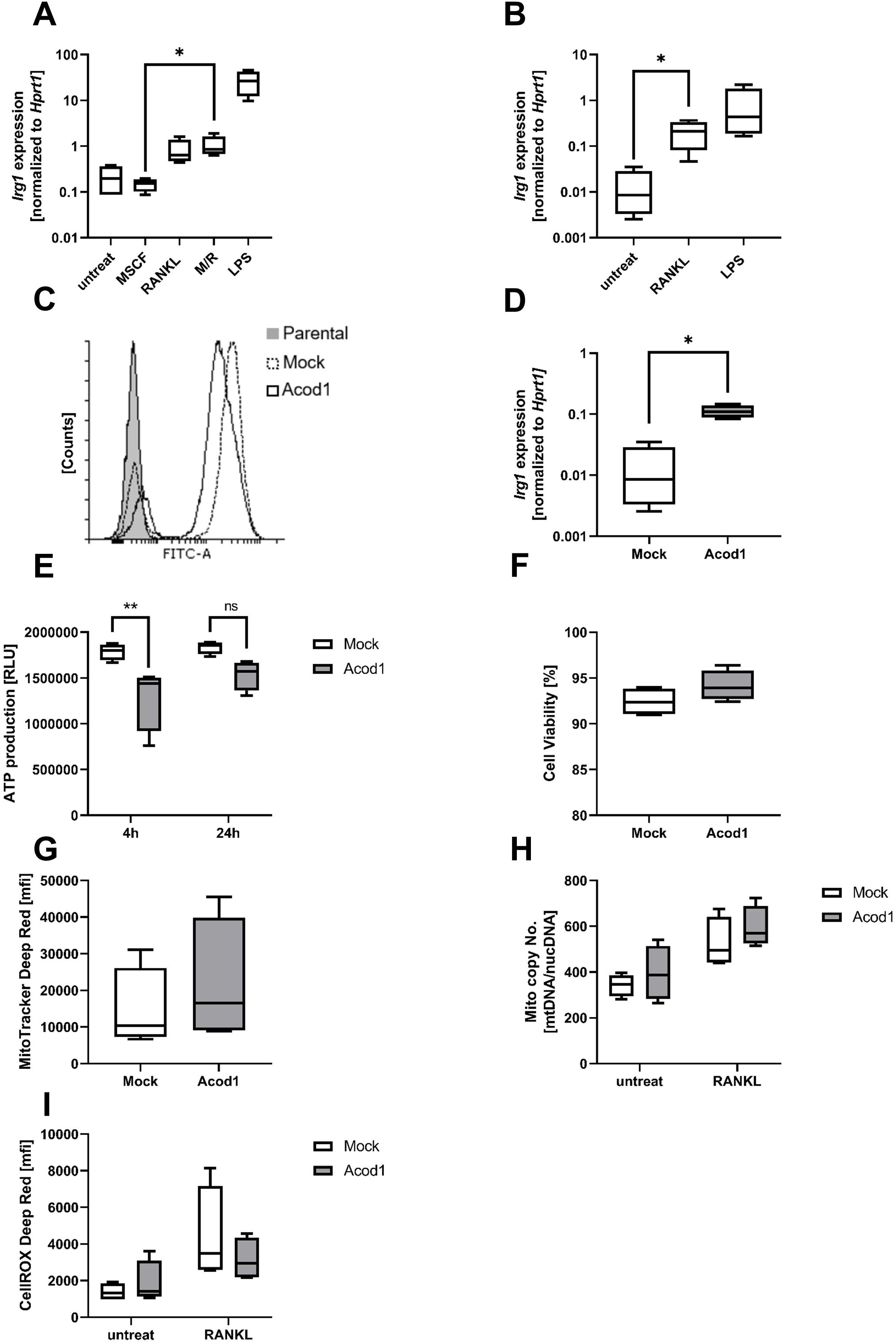
RANKL induces *Irg1* expression. **(A,B)** *Irg1* gene expression analysis by qPCR. BMDMs **(A)** and RAW 264.7 macrophages **(B)** were stimulated with 10 ng/mL LPS and 50 ng/mL RANKL for 6 h, respectively. BMDMs were additionally treated with 25 ng/mL M-CSF. Relative *Irg1* expression levels were normalized to the reference gene *Hprt1*(n=4). **(C)** FACS analysis of lentiviral-transduced Mock and Acod1 overexpression cells for eGFP fluorescence. Parental RAW264.7 macrophages were included as a negative control. **(D)** *Irg1* gene expression analysis of Mock and Acod1 overexpressing cells by qPCR. Relative *Irg1* expression levels were normalized to reference gene *Hprt1* (n=4) **(E)** The ATP content in Mock and Acod1 overexpressing cells was assessed by Cell TiterGlo^®^ Assay analysis after 4 or 24 h cultivation (n=4). **(F)** Cell viability of Mock and Acod1 cells was assessed by Zombie NIR™ staining. The percentage of live cells was shown. **(G)** Mitochondrial activity was evaluated by MitoTracker™ Deep Red. Both Mock and Acod1 cells were cultured for 24 h (n=4). **(H)** The mitochondrial DNA content was assessed by determining the mitochondrial copy number. Both Mock and Acod1 cells were stimulated with 50 ng/mL RANKL for four days. The number of mitochondria per cell were quantified by qPCR. Mitochondrial DNA (mtDNA) copies were normalized with nuclear DNA (nucDNA) (n=4). **(I)** Cellular oxidative stress was measured by CellROX™ Deep Red reagent. Mock and Acod1 cells were stimulated with RANKL. Cells were evaluated by FACS on day 3 after 30 min CellROX™ staining (n=4). Statistical analysis was performed using Mann-Whitney test.

During osteoclast differentiation, mitochondrial biogenesis is activated to support the increased energy requirement. As the TCA cycle activity is affected in Acod1 overexpressing cells, we quantified mitochondrial activity using Mitotracker Deep Red stain. We found the mitochondrial activity to be increased in Acod1 cells (Figure 1G). As differentiating osteoclasts are known to undergo mitochondrial biogenesis, we then investigated the mitochondrial copy number to see if compensatory mechanisms were activated. As expected, RANKL treatment caused an upregulation of the mitochondrial copy number and this effect was more pronounced in Acod1 cells, however, not reaching statistical significance. Here, the upregulation was detectable already under untreated conditions, although to a lower extent (Figure 1H). We therefore suggest that the increase in mitochondrial activity is primarily caused by an increase in mitochondrial mass. Therefore, we next determined cellular ROS production by staining the cells with the CellRox^®^ Deep Red reagent (Figure 1 I). While there was no difference between untreated cells, RANKL treatment caused an increase in ROS in both cells as ROS is an important secondary messenger that supports osteoclast formation. Acod1 cells, however, showed a clear decrease in ROS compared to RANKL-treated Mock cells, presumably due to a decrease in reverse electron transfer (RET) as the consequence of SDH inhibition.

### 3.2 Acod1 cells show reduced osteoclast formation

As ROS plays important role in osteoclast formation, we investigated the ability of Mock and Acod1 cells to differentiate with RANKL into multi-nucleated, tartrate resistant acidic phosphatase (TRAP)- positive osteoclasts. On day 5 of differentiation, cells were stained and counted. Indeed, for Acod1, the number of osteoclasts was significantly reduced, although there was not a complete abrogation of osteoclast formation (Figure 2A). To investigate if this reduction was caused by a difference in kinetics, we compared osteoclast differentiation on days five and six to see if Acod1 cells might catch up at a later time-point (Figure S2). Longer incubation times indeed showed that there was a delay in differentiation. While on day six the number of Mock cell-derived osteoclasts already decreased due to apoptosis, Acod1 osteoclasts still fused and generated more osteoclasts. However, the number was still lower than that of day five osteoclasts. Looking at the cell morphology (Figure 2B), it became clear that Acod1 osteoclasts stained less intensive for TRAP although the size did not seem to differ significantly. Next, we analysed the expression of osteoclast specific genes during differentiation (Figure 2C-J). *Nfatc1* that gets auto-amplified during the initial phase of osteoclast differentiation, only showed a significant difference between Mock and Acod1 cells late during differentiation (day3). *Dcstamp*, a marker for osteoclast fusion remained unchanged, but all other genes were severely suppressed in Acod1 cells (Figure 2D-J). The investigated genes included *Acp5, Atp6vod2, Ocstamp*, *Calcr*, *Oscar*, and *Ctsk*.

**Figure 2:**
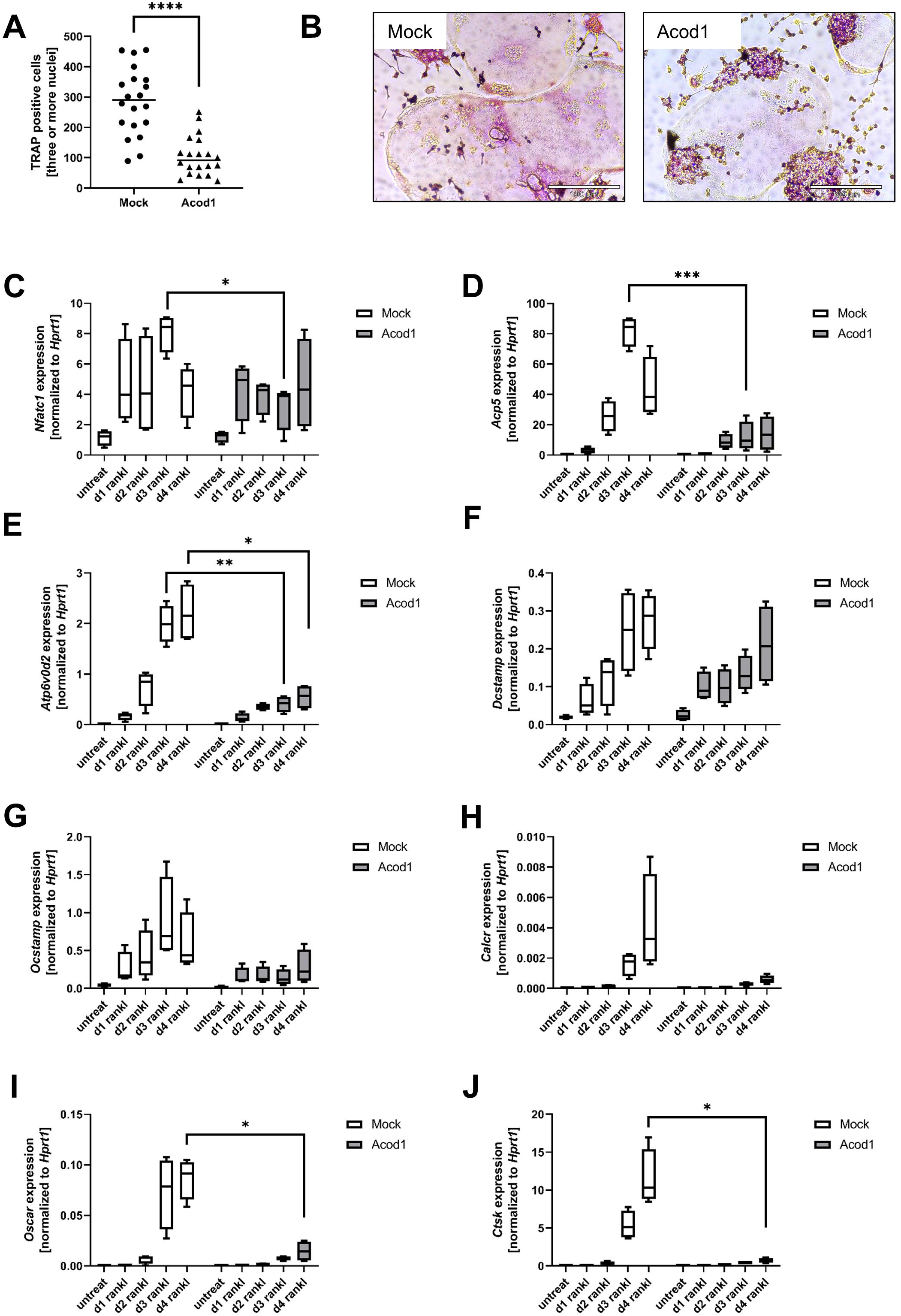
Acod1 suppresses osteoclast differentiation. **(A)** Quantification of TRAP staining; **(B)** Representative pictures of TRAP staining. Both Mock and Acod1 cells were seeded as duplicates and stimulated with 50 ng/mL RANKL. TRAP stainings were performed after five days of differentiation (n=10). **(C-J)** Osteoclast differentiation analysis by qPCR. Gene expression levels of several characteristic osteoclast genes (*Nfatc1, Acp5, Atp6vod2, Dcstamp, Ocstamp, Calcr, Oscar, Ctsk*) were evaluated by qPCR over a differentiation time course. Samples were harvested at the indicated time-points. The relative gene expression levels were normalized to reference gene *Hprt1*. (n=4) Statistical analysis was performed using a Mann–Whitney test.

### 3.3 Activation of RANKL-induced signaling pathways

Next, we verified the expression of RANK at the cell surface to rule out that Acod1 causes an inhibition of RANK surface expression. The FACS data show that RANK localisation of Acod1 cells is reduced by app. 25%, however, this does not explain the absence of NFATc1-mediated osteoclast gene induction (Figure 3A). As ROS levels as well as osteoclast gene expression were decreased, resulting in reduced osteoclastogenesis, we expected a reduction of RANKL-mediated NFATc1 activation since NFATc1 is the master regulator of osteoclastogenesis. To address this, we determined the activation of NFATc1, NFATc2, the activated NF-κB subunit phospho-p65, as well as BLIMP1, a transcriptional repressor that counter-regulates the Bcl6-induced repression of osteoclast genes. Unexpectedly, whole cell lysates showed that neither NFATc1, NFATc2, p-p65 or BLIMP1 were downregulated in Acod1 cells on day 0. Instead, we saw a pronounced upregulation of protein expression or activity in untreated Acod1 cells. This increase was also visible during differentiation, though not as pronounced as in the untreated state (Figure 3B, 3C). This suggests that the observed basal NFATc1 expression does not translate in the upregulation of the osteoclast differentiation program. Therefore, we next checked if NFATc1 was perhaps impaired in its ability to translocate to the nucleus. Surprisingly, we found NFATc1 to be present in the nucleus in untreated Acod1 cells (Figure 3D). Mock cells showed the expected translocation of NFATc1 to the nucleus only after 6 hours of RANKL treatment, while there was no further increase in NFATc1 expression in Acod1 cells treated with RANKL. We therefore investigated if nuclear location was not sufficient to trigger NFAT transcriptional activity using an NFAT luciferase reporter assay (Figure 3E). Interestingly, Acod1 cells did show residual NFAT activity in the absence of treatment which was not observed for Mock cells. In summary, these data suggest that Acod1 overexpression causes a constitutive induction of NFATc1 expression, nuclear translocation and transcriptional activity. While the missing responsiveness to RANKL observed in the western blot corresponds with the observed phenotype of Acod1 cells, it did not explain the constitutive activation of NFATc1 in the untreated state.

**Figure 3:**
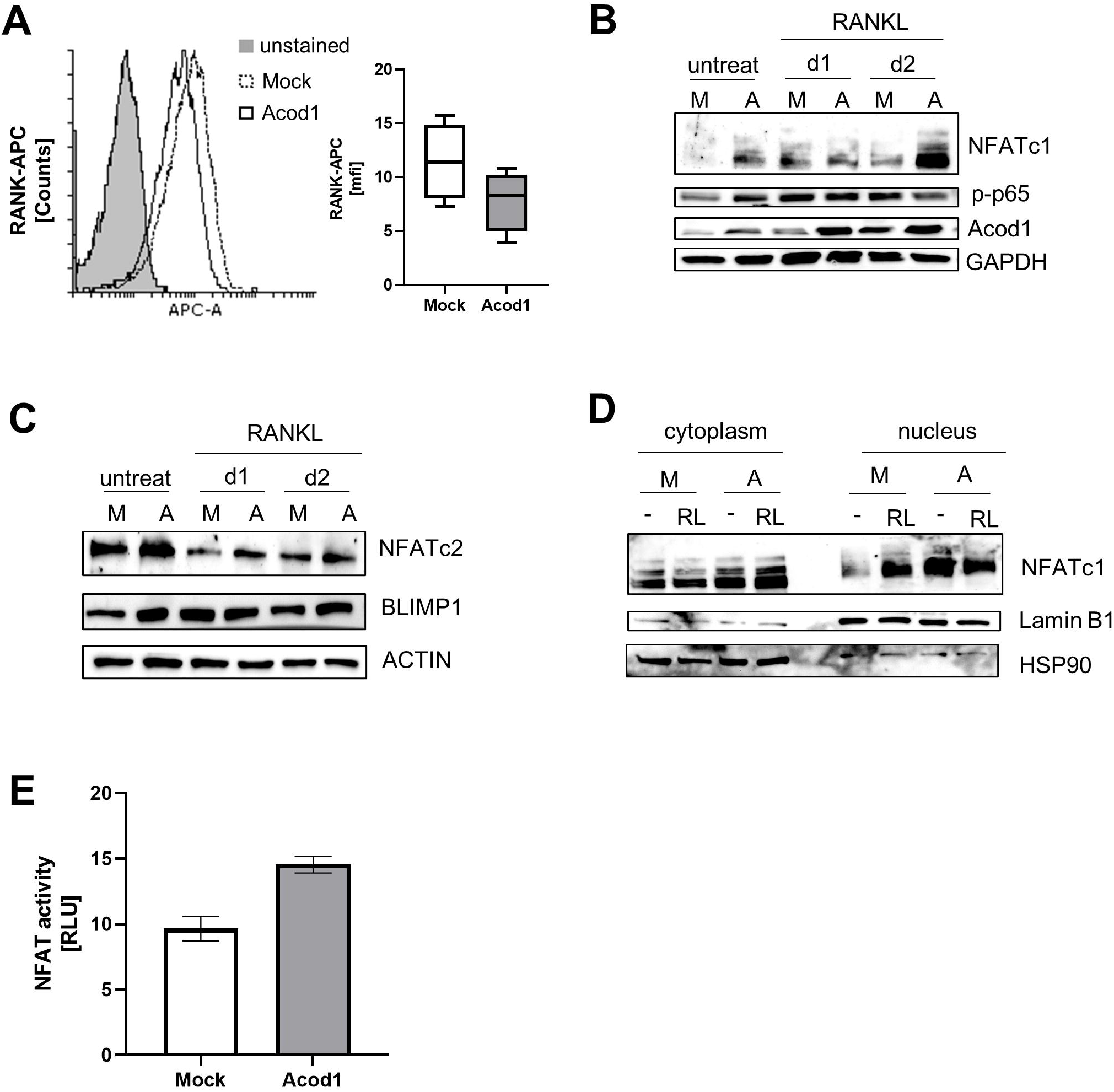
Activation of RANKL-induced signaling pathways. **A)** RANK surface expression was measured by FACS 1 day after seeding using an allophycocyanin (APC)-conjugated antibody. An unstained sample was included as control. The histogram shows one representative example and the box plot summarises all experiments (n=4) **(B,C)** Western Blot analysis of the osteoclast differentiation associated transcriptional factors NFATc1, phospho-p65 **(B)**, NFATc2, and Blimp1 **(C)**. Mock and Acod1 cells were stimulated with 50 ng/mL RANKL and then harvested at the indicated time-points. GAPDH and Actin were loaded as control. One typical example of 2 replicated experiments is shown. **(D)** Subcellular location analysis of NFATc1 by Western Blot. Nuclear extraction was performed after 6 hours of 50 ng/mL RANKL stimulation. Lamin B1 and HSP90 were loaded as nuclear and cytoplasmic control, respectively. **(E)** Transcriptional activity of NFAT was analysed by luciferase assay. Transient transfection with the reporter constructs was performed after seeding. Cells were lysed and evaluated after 6 hours of stimulation. The relative luminescence units were normalized to the *Renilla* luciferase activity. Technical duplicates of one representative experiment are presented (n=4).

### 3.4 Succinate receptor GPR91 expression and activity is regulated by itaconate

Next, we investigated if Acod1 overexpression might impact other metabolites, such as lactate and succinate. They are known to be important for macrophage function and osteoclast differentiation, respectively (7), and could represent the link to the observed missing NFATc1 responsiveness to RANKL. We therefore determined the gene expression of succinate dehydrogenase of the complex II in the electron transport chain (*Sdha*), as well as lactate dehydrogenase (*Ldha*) and succinate receptor (*Gpr91*) in untreated cells and during differentiation with RANKL (d1-d5) and compared between Mock and Acod1 cells. Figure 4A shows that the expression of *Sdha* was not regulated during the course of differentiation. In addition, there was no significant difference between Mock and Acod1 cells, although *Sdha* expression of was slightly decreased in Acod1 cells (Figure 4A). For *Ldha* we observed an upregulation at later time-points during differentiation, however, this was observed for both cell lines (Figure 4B). As itaconate causes accumulation and export of succinate, we also investigated the gene expression of the succinate receptor GPR91. Figure 4C shows that in both cell lines, *Gpr91* transcription is not induced prior to RANKL treatment. Stimulation with RANKL strongly induced *Gpr91* expression, starting at day 2 in Mock cells. In Acod1 cells this effect was similar but much lower and the expression was reproducibly reduced from day 2-5 compared to Mock control cells. Bioinformatical analysis of the *Gpr91* promoter region showed that it contains multiple NFAT binding sites, suggesting that the receptor is a direct target of RANKL-mediated NFAT activation (Figure S3). Next, we investigated whether the decrease in gene transcription was a direct effect of itaconate. Stimulation of Mock cells with different concentrations of octyl-itaconate ranging from 50-200 μM showed that octyl-itaconate dose-dependently inhibited *Gpr91* expression (Figure 4D). We then determined the localisation of GPR91 at the cell surface to see if the results observed for *Gpr91* expression were reflected by an inhibition of GPR91 surface localisation as well. However, we found that GPR91 is expressed already at high levels in untreated Mock cells (macrophage state) and that the receptor surface expression decreases during osteoclastogenesis. Acod1 cells showed significantly reduced levels in the untreated state (app. 45%), but the receptor was still abundantly expressed at the cell surface. The expression was further reduced during RANKL treatment for both cell lines (Figure 4E). To test if the amount of GPR91 at the surface of Acod1 cells was sufficient to induce succinate-induced signalling, we treated the cells with 500 μM succinate for 6 hours and quantified NFAT transcriptional activity in a luciferase assay (Figure 4F). The data demonstrate that GPR91 on Acod1 cells is still able to trigger NFAT activation through its associated heterotrimeric G protein Gαq in response to succinate treatment. This suggests that the observed residual NFAT activity can be caused by succinate-mediated signalling which accumulates in Acod1 cells as a consequence of SDH suppression (13).

**Figure 4:**
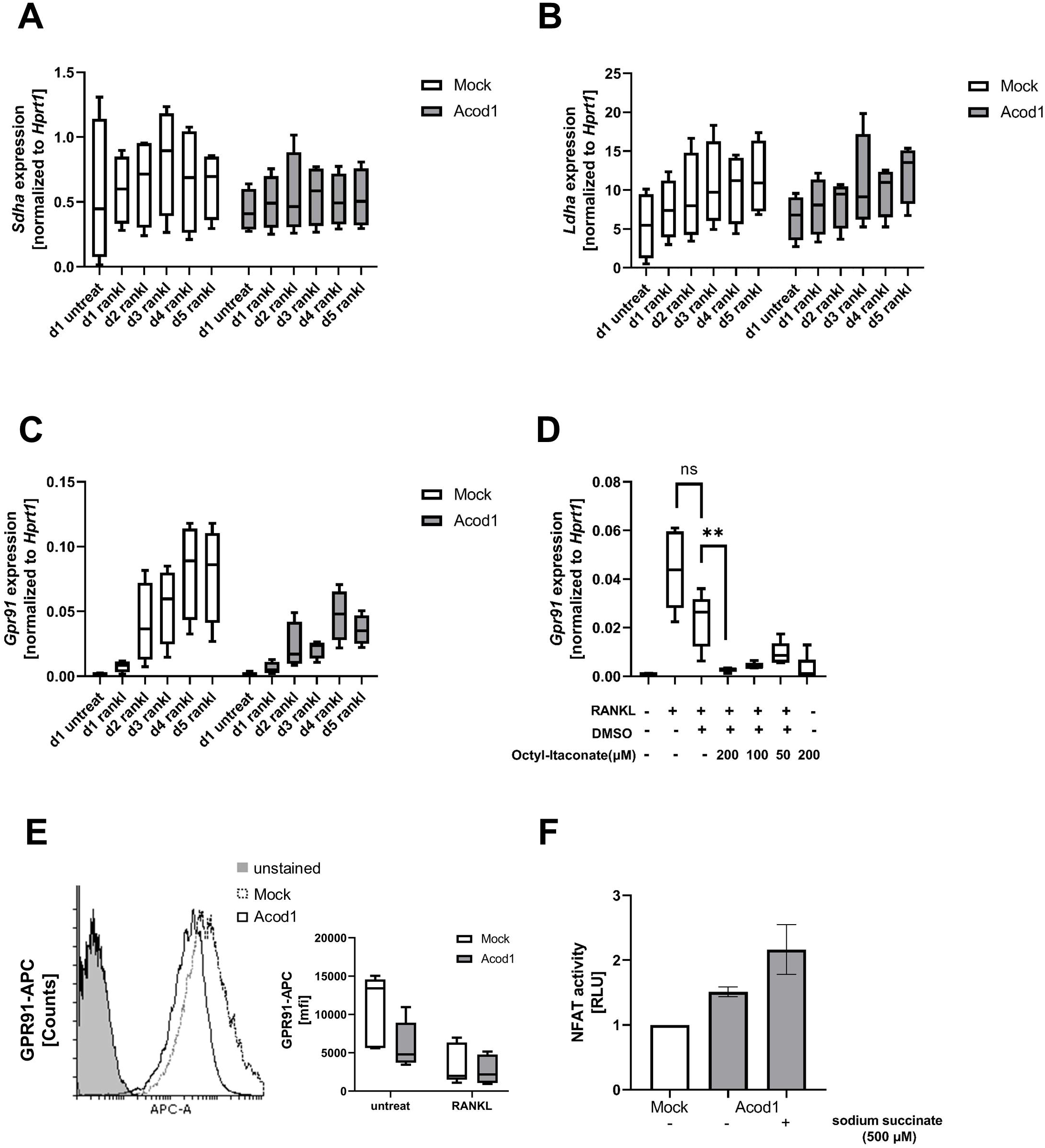
Succinate receptor GPR91 expression and activity is regulated by itaconate. **(A-C)** Gene expression analysis of *Sdha* **(A)**, *Ldha* **(B)** and *Gpr91***(C)**. Cells were stimulated with 50 ng/mL RANKL and harvested as the indicated time-points. The relative gene expression was normalized to the reference gene *Hprt1* (n=4). **(D)** *Gpr91* expression analysis with co-stimulation of RANKL and itaconate. Cells were seeded and stimulated with 4-octyl itaconate 3 h prior to 50 ng/mL RANKL stimulation. On day 3, the cells were re-stimulated with itaconate and RANKL. Sample were harvested on day five. A solvent control was included to control for DMSO-mediated effects on cell differentiation and gene expression. The relative gene expression was normalized to the reference gene *Hprt1* (n=4). **(E)** GPR91 surface expression was measured by FACS after 3 days of 50 ng/mL RANKL stimulation. An allophycocyanin (APC)-conjugated antibody was applied. The histogram shows one representative example that compares GPR91 surface localisation of untreated cells and the box plot summarises the experiments for untreated and RANKL-treated conditions (n=4).

### 3.5 Acod1-deficient mice do not display altered OC formation

To investigate whether the absence of Acod1 would influence osteoclastogenesis and ultimately affect bone formation, we used primary bone marrow-derived macrophages from Acod1-deficient mice (Acod1^em1(IMPC)J^/J mice). When we investigated the ability of these cells to differentiate into osteoclasts, we observed a small, statistically not significant increase in the number of TRAP positive, multinuclear cells (Figure 5A). Interestingly, TRAP staining of osteoclast revealed a more intense staining of the osteoclasts for Acod1-deficient cells (Figure 5B), while we had seen a reduced staining in Acod1 overexpressing RAW264.7 macrophages. When we looked at the expression of osteoclast-specific genes we found again that *Nfatc1* expression was not changed between wildtype and knock-out mice (Figure 5C). However, the expression of *Acp5, Ctsk* and *Atp6vod2*, which had been the most significantly reduced genes in Acod1 overexpressing cells, showed a comparable expression kinetics with a tendency of higher induction in the Acod1-deficient cells. However, none of this was statistically significant. In addition, macrophages from Acod1-deficient mice showed no changes in viability, no significant changes in ATP production and a higher induction of RANKL-mediated NFATc1 activation in Western Blot analysis (Figure S4).

**Figure 5:**
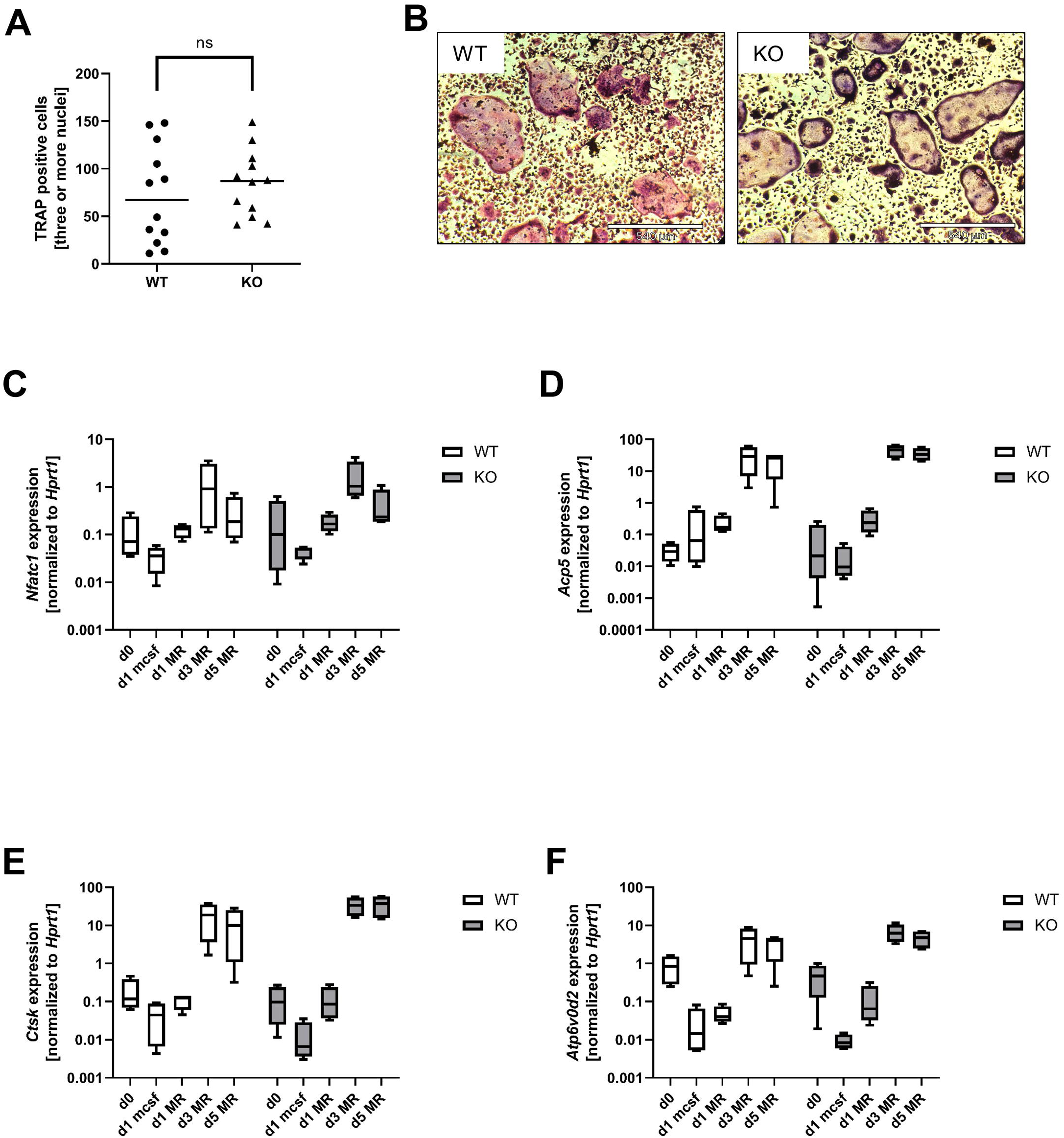
Acod1-deficient mice do not display altered OC formation. **(A)** Quantification of TRAP staining; **(B)** Representative pictures of TRAP staining; Wildtype and *Acod1^-/-^* BMDMs were seeded as duplicates and stimulated with 50 ng/mL RANKL and 25 ng/mL M-CSF. TRAP staining was performed after five days of differentiation (n=6). **(C-F)** Analysis of the osteoclast genes expression by qPCR. Gene expression levels of several characteristic osteoclast genes (*Nfatc1, Acp5, Ctsk, Atp6vod2*) were evaluated by qPCR over a differentiation time course. Samples were harvested at the indicated time-points. The relative gene expression levels were normalized to reference gene *Hprt1* (n=4).

### 3.6 Bone phenotype and arthritis

To evaluate if the observed small differences might translate into a change in bone morphology, we investigated the composition of the femur and tibia. An increase in osteoclastogenesis and therefore in bone degrading cells, would result in the loss of bone and in the decrease of bone density. To evaluate the differences in the bone phenotype of WT and Acod1-deficient mice, μCT scans were performed. After μCT scanning, the VOIs were selected in the resulting radiographs (Figure 6A) and were analysed concerning the following parameters: Total volume (TV), bone volume (BV), bone surface (BS) and mean greyscale index, which is a measure for bone density. While the μCT analysis did not reveal any differences in bone mass (BV/TV) of the WT and Acod1-deficient samples (Figure 6B), alterations in bone compactness (BS/BV) and density were observable (Figure 6C and 6D). Both, tibiae and femora of the Acod1-deficient mice showed higher BS/BV values, thus a lower bone compactness compared to the WT mice (Figure 6C). Accordingly, the bone density was slightly decreased in the femora and tibiae of ACOD-1 ko mice as indicated by a lower mean grey value (Figure 6D). The differences were statistically significant, but seem to be rather marginal. Therefore, we conclude that at least under healthy conditions there are no physiologically relevant changes of the bone phenotype between wildtype and knock-out mice. This suggests that the absence of Acod1 has a lower impact than the accumulation of Acod1-produced itaconate with respect to osteoclastogenesis.

**Figure 6.**
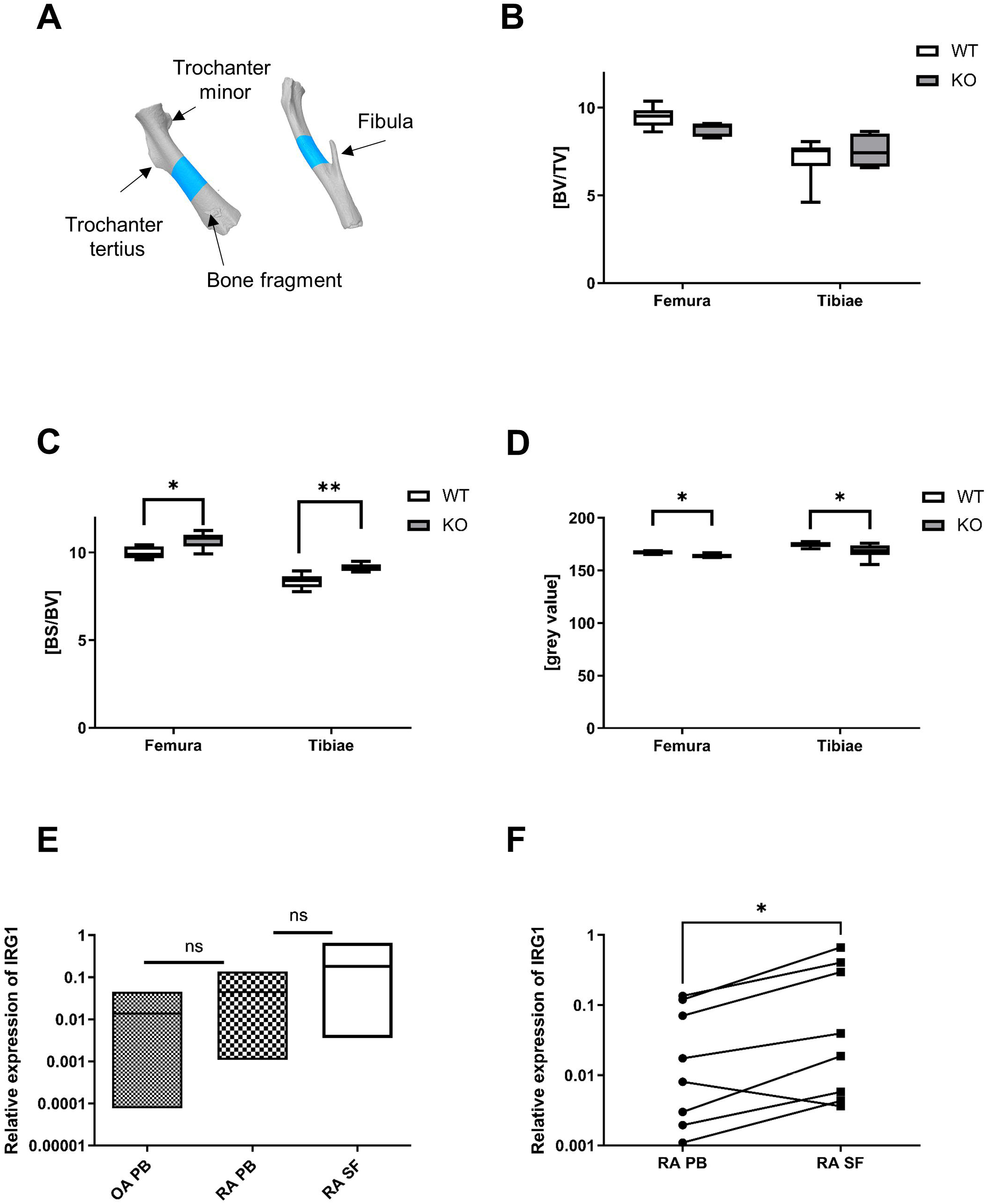
μCT analysis of murine femora and tibiae reveals only slight differences in bone size, structure and density comparing wildtype and *Acod1^-/-^* mice. **A)** 3D images generated from the μCT scans with CTAn and CTVol. The blue fragments show the selected VOIs for the analysis. Left: Example of a femur analysis. The VOI was selected distal from the Trochanter tertius. Right: Example of a tibia analysis. The VOI was selected proximal from the insertion of the fibula. In both cases, 150 images were selected as the VOI. **(B-D)** Results of the final analyses of the bone samples. BV/TV is a measure for the bone volume in the selected VOI **(B)**. BS/BV describes the compactness of the bone. If the bone surface (BS) is increasing in comparison to the BV, the analysed sample compactness decreases **(C)**. The grey value is the mean greyscale index and displays the bone density (D). The numerical results can be found in supplementary table 1. n = 6 (2 bones from 3 mice each), Mann-Whitney U, ns = non-significant, * p < 0.05, ** p < 0.005 **(E,F)** The relative expression of *Irg1* was measured in cells derived from peripheral blood (PB) and autologous Synovial fluid (SF) of RA patients with active disease; peripheral blood of OA patients using quantitative real time pCR **(E)**. Patient samples from OA, a degenerative joint disease, were considered as a study control for RA, a chronic inflammatory joint disease. The graphs represent the relative expression of the *Irg1* in OA and RA samples (mean ± SD; n = 8) **(E)** and the comparison of *Irg1* PB/SF values for each patient **(F)**. Statistical analysis was performed using a non-parametric t-test.

There are conflicting data regarding the association between the levels of itaconate and disease progression in RA patients. Therefore, we performed a small pilot study and checked *Irg1* in cells derived from peripheral blood (PB) and synovial fluid (SF) of RA patients, to understand if there is any difference in the expression of *Irg1* at the site of inflammation compared to the periphery. In addition, we analysed *Irg1* in cells derived from PB of osteoarthritis (OA) patients as a non-inflammatory disease control (Figure 6E). Overall, we found a statistically significant increase in *Irg1* expression in SF-derived monocytes compared to PB in RA patients (Figure 6F). The increase in *Irg1* expression could suggest that *Irg1* might rather be a marker of resolution than inflammation as it shifts pro-inflammatory macrophages to a less activated state (31) and limits osteoclastogenesis as suggested by the data obtained in our *in vitro* model.

## 4. Discussion

Both, Irg1 and GPR91 were found in the 1990s, however it took until 2004 to deorphanize GPR91 as the succinate receptor (33) and until 2013 to identify the enzymatic function of *Irg1* as the aconitate decarboxylase (12). Since then, a wealth of new information on the influence of metabolism on the regulation of innate and adaptive immune responses has been obtained. The immunomodulatory function of itaconate has been reviewed in a number of publications and its central functions comprise the inhibition of SDH, the NLRP3 inflammasome and glycolysis in addition to its resolving roles due to activation of Nrf2, a central transcription factor in the regulation of anti-oxidants genes and ATF3, an anti-inflammatory transcription factor, especially in dendritic cells and macrophages (34). As bone-resorbing osteoclasts develop from the monocyte/macrophage lineage, we studied the effect of Acod1 on the differentiation of osteoclasts. Here, we used an overexpression system to see the endogenous effects of itaconate as the different chemical properties of esterized compounds, such as dimethyl itaconate or 4-octylitaconate, might result into different biological activities. Although the upregulation of *Irg1* expression in response to RANKL treatment suggested that its expression might be necessary for osteoclastogenesis, Acod1 overexpressing cells expressed a significantly reduced number of osteoclasts demonstrating that itaconate is able to limit the extent of osteoclastogenesis. This is in line with results from Sun and colleagues who showed in 2019 that addition of octyl itaconate to developing osteoclasts inhibited osteoclastogenesis through the activation of ROS-quenching Nrf2 signaling in an osteoporosis model of ovariectomised mice (32). We can corroborate their finding that ROS production was supressed and that also endogenously produced itaconate prevents osteoclastogenesis. In addition, we also investigated whether the absence of itaconate might enhance osteoclastogenesis but this does not seem to be the case, at least in young and healthy mice, as Acod1-deficient animals did not display a distinct bone phenotype.

Our model system suggests that in addition to the observed decreased levels of ROS, itaconate acts via regulation of succinate-mediated signalling. As the inhibition of SDH activity causes an accumulation and subsequent secretion of succinate, succinate can extracellularly activate its cognate receptor GPR91 (13). In rodents and humans, GPR91 is expressed in various tissues and cells types and also in myeloid cells such as DCs and macrophages as well as T and B cells of the adaptive immune system [reviewed in (35)]. Upon ligand binding, GPR91 can activate Gαi and Gαq signalling and the preference for these pathways seems to be cell type specific. In macrophages, Gαq-mediated Ca^2+^ release and subsequent NFATc1 activation is the predominant pathway (36). Our data show that in addition to NFATc1 expression, nuclear translocation and transcriptional activity, Acod1 overexpression also activates NF-kB (p-p65), and enhances AP-1 (c-jun) and BLIMP1 expression. While NF-kB and AP-1 are known targets of Gαq, there are no data on the regulation of the transcriptional repressor BLIMP1 by GPCR signalling, although there might be a connection between BLIMP1-mediated differentiation of B cells into plasma cells and the activation of signalling via the GPCR sphingosine receptor 1 (37).

Initially it was assumed that in healthy cells, GPR91 must be quiescent as GPR91 activation is indeed best known for its pathophysiological consequences, such as cardiac hypertrophy, ischemic retinopathy diabetic nephropathy and the overactivation of macrophages in RA or in cancer (20). Here, succinate functions as a danger associated molecular pattern (DAMP) signal that helps the body to sense changes in metabolic activity (38). The few physiological functions include retinal vascularisation and the stimulation of blood pressure in the kidney through activation of the renin-angiotensin system, but an increase of extracellular succinate can also be observed after physical exercise and food ingestion (38–41). Here, succinate is released from the exercising myofibers and acts as a so-called “metabokine” in a paracrine manner to initiate muscle extracellular matrix remodelling which helps skeletal muscles to adapt to exercise training (39). In the context of macrophage activation, succinate can trigger both, an anti-inflammatory or a pro-inflammatory response, depending on the context. GPR91 shifts tumour-associated macrophages into a phenotype that dampens the immune response and recent data from Trauelsen et al. also suggest that succinate is anti-inflammatory in nature and strengthens the phenotype of alternatively activated M2 macrophages (36). In RA, however, macrophages and osteoclasts show a pro-inflammatory phenotype in response to succinate and in such pathologies, significantly increased levels of circulating succinate can be found (42, 43). These contradictory findings imply that succinate can reinforce existing signals transmitted by the microenvironment, and therefore act either pro- or antiinflammatory (44). In the absence of such a signal as in our untreated Acod1 overexpressing cells, succinate itself might not trigger any measurable changes in cell activation or regulation as a second stimulus is missing. Indeed, we did not observe major changes in cytokine production in Acod1 overexpressing cells (data not shown). A similar scenario was found for a bacterial protein toxin from *Pasteurella multocida* (PMT) that constitutively activates Gαq. PMT additionally triggers the production of pro-inflammatory cytokines through NF-kB signalling, and only the simultaneous activation of both pathways allowed for RANKL-independent osteoclastogenesis (25). Similarly, succinate and GPR91 stimulate osteoclastogenesis in the context of type II diabetes, where there is an underlying inflammation with the production of pro-inflammatory cytokines (21). Our model will be helpful to understand if GPR91 actually targets specific genes in quiescent cells. To define the phenotype of succinate-activated Acod1 cells, ChIP-seq analysis with NFATc1 will help to understand the expression of succinate-activated, NFAT-induced genes. It is also plausible that the increased amounts of metabolites prevent chromatin accessibility, e.g. through succinylation. Results on the expression of histone mRNA remained inconclusive (Figure S5), but points to differences in the regulation of chromatin structures, especially at later time-points, when Acod1 cells seem to respond to RANKL more efficiently.

Or data discovered an interesting discrepancy between *Gpr91* gene expression and localisation of the protein at the cell surface. First of all, our data suggest that GPR91 is a RANKL-inducible gene, as GPR91 contains several NFAT binding motifs in its reporter region (Figure S3), which explains the observed RANKL-dependent upregulation on the mRNA level. However, the receptor is additionally and more importantly regulated at the protein level. While untreated Mock cells show a high expression of GPR91 at the cell surface, the gene is not induced under these conditions. Vice versa, the increase in *Gpr91* mRNA during differentiation goes along with a remarkable downregulation of its localisation at the plasma membrane, but the reason for this regulation requires in-depth investigation. Arrestins are known regulators of GPCRs which support desensitisation and internalisation by tagging activated GPCRs towards clathrin-coated vesicles and endocytosis. There are also examples of GPCRs that trigger ubiquitination which causes endocytosis (45). Information on the regulation of GPR91 surface localisation is currently missing, but Robben et al. reported earlier that GPR91 gets desensitized in the absence of receptor internalization (38, 46). Alternatively, it is possible that in the presence of increased levels of itaconate, the receptor is not transported correctly to the cell surface, for examples due to a change in glycosylation which has been reported for other GPCRs (47). As many publications refer to mRNA levels when talking about GPR91 expression, these data have to be evaluated cautiously and need to be complemented with protein expression data specifically looking at its the localisation at the plasma membrane.

The recent increase in our understanding of immunometabolism holds the promise of new approaches to treat inflammation-induced bone loss, which occurs in a number of diseases, such as RA, osteomyelitis or periodontitis. As these diseases go along with increased osteoclast formation and enhanced glycolysis, pharmacological approaches aiming at shifting the cellular metabolism towards the TCA cycle are currently being discussed. However, *in vivo* models of auto-inflammatory diseases mostly address the possibility to manipulate aberrant T cell activity and the effect of macrophage/osteoclast plasticity is less well understood. Recently, Taubmann et al. reported on an approach of osteoclast reprogramming during the treatment of osteoporosis, where inhibition of glycolysis did not interfere with the process of differentiation but effectively decreased the resorptive capacity of the osteoclasts (48). Regarding itaconate, a recent study detected a substantial amount of the metabolite in plasma samples collected from rheumatoid arthritis (RA) patients, but the level of itaconate after 3 months of treatment negatively correlated with the disease activity score (DAS28), suggesting that itaconate acts as an anti-inflammatory marker that could be a read-out for improved resolution of the inflammation (49). This finding correlates with our *in vitro* data, where the overexpression Acod1 diminishes osteoclastogenesis, and the observed increase in *Irg1* between the SF and PB samples of RA patients. A murine transgenic arthritis model based on the overexpression of TNF-α was discussed to be in conflict with this view, as the metabolic profiling by mass spectrometry showed elevated itaconate concentrations in synovial fibroblasts from Tg197 transgenic mice that reversed upon TNF-α inhibitor treatment (50). However, as TNF-α is a strong inducer of *Irg1* (30), it is clear that overexpression of the cytokine will result in increased *Irg1* and itaconate levels, while blocking TNF-α will reverse this. Enhanced succinate levels which can be a consequence of itaconate-mediated inhibition of SDH activity, are detected in RA patients where succinate contributes to enhanced osteoclastogenesis (21, 22). Succinate receptor deficiency on the other hand attenuates arthritis, as it reduces the migration of DCs into the lymph nodes and thus prevents the expansion of auto-inflammatory Th17 cells (51). Our data therefore support the idea that *Irg1* can be regarded as a marker for resolution as itaconate limits *Gpr91* expression and limits succinate-mediated GPR91 signalling and NFAT activation by decreasing the availability of the receptor. While our data are purely *in vitro*, we hope to contribute to a better understanding of the observed discrepancies and add to the understanding of new layers of regulation in the crosstalk between itaconate, succinate and succinate receptor and their possible targeting for immune modulation.

## 6. Author Contributions

YG, FVK and SC performed experiments and collected data. ES, DS, JS and WW analysed data. KFK conceptualised and designed the study and wrote the manuscript. All authors contributed to manuscript revision, read, and approved the submitted version.

## 8. Acknowledgments

We are grateful for the support of Lavinia Flaskamp (Heidelberg), who helped to start the project, Gabriele Sonnenmoser for technical assistance, Friederike Berberich-Siebelt (Würzburg) for helpful discussions and Daniel Mertens (Ulm) for the generous gift of the NFAT luciferase construct.

## 10. Supplementary Material

Supplementary Material will be uploaded separately.

