## Supplementary files for "Acod1 negatively impacts osteoclastogenesis via GPR91-mediated NFATc1 activation"

### Supplementary Material

#### 1 Supplementary Methods

##### Transcription factor binding site prediction

*Mus musculus Sucnr1* genomic sequence (Gene ID: 84112) was used to predict putative transcription factor binding sites (TFBS). The genomic data was analysed in R/Bioconductor (1) environment utilising the package TFBSTools (2) based on the latest JASPAR 2020 database (3) of curated, non-redundant transcription factor binding profiles. All predicted NFAT TFBS candidates with both a minimal score of 0.8 and a p-value below 0.001 were visualised with the Gviz package (4) as overlay to the ideogram track of *Mus musculus* UCSC (mm39) reference genome (https://www.bioconductor.org/packages/release/data/annotation/html/TxDb.Mmusculus.UCSC.mm39.refGene.html).


1. Team RC. R: A language and environment for statistical computing. R Foundation for Statistical Computing. Vienna, Austria2021.

2. Tan G, Lenhard B. TFBSTools: an R/bioconductor package for transcription factor binding site analysis. Bioinformatics. 2016;32(10):1555-6.

3. Fornes O, Castro-Mondragon JA, Khan A, van der Lee R, Zhang X, Richmond PA, et al. JASPAR 2020: update of the open-access database of transcription factor binding profiles. Nucleic Acids Res. 2020;48(D1):D87-D92.

4. Hahne F, Ivanek R. Visualizing Genomic Data Using Gviz and Bioconductor. Methods Mol Biol. 2016;1418:335-51.


#### 2 Supplementary Tables and Figures

##### 2.1 Primers used in RT-PCR:

| | |
|---|---|
| *Irg1* | Forward: 5'-CAGCTCTATCGGAAGCCCTG-3'<br>Reverse: 5'-CAGAAACTTGGACGCAGCAG-3' |
| *Hprt1* | Forward: 5'-GGGGACATAAAAGTTATTGGTGG-3'<br>Reverse: 5'-CATTTTGGGGCTGTACTGCT-3' |
| *Nfatc1* | Forward: 5'-CAGGGCTCACTATGAGACGG-3'<br>Reverse: 5'-AGCTGTAGCGTGAGAGGT-3' |
| *Acp5* | Forward: 5'-TTCCAGGAGACCTTTGAGGA-3'<br>Reverse: 5'-GGTAGTAAGGGCTGGGGAAG-3 |
| *Atp6v0d2* | Forward: 5'-TCAGATCTCTTCAAGGCTGTGCTG-3'<br>Reverse: 5'-GTGCCAAATGAGTTCAGAGTGATG-3' |
| *Dcstamp* | Forward: 5'-AAAACCCTTGGGCTGTTCTT-3'<br>Reverse: 5'-GTTCCTTGCTTCTCTCCACG-3' |
| *Ocstamp* | Forward: 5'-TGGGCCTCCATATGACCTCGAGTAG-3' |

| | |
|---|---|
| | Reverse: 5’-TCAAAGGCTTGTAAATTGGAGGAGT-3’ |
| *Calcr* | Forward: 5’-AGAGTGAAAAGGCGGAATCT-3’<br>Reverse: 5’-TTTGTACTGAGCATCCAGCA-3’ |
| *Oscar* | Forward: 5’-AGGGAAACCTCATCCGTTTG-3’<br>Reverse: 5′-GAGCCGGAAATAAGGCACAG-3′ |
| *Ctsk* | Forward: 5’-AGGGAAGCAAGCACTGGATA-3’<br>Reverse: 5’-GCTGGCTGGAATCACATCTT-3′ |
| *Gpr91* | Forward: 5’-GACAGAAGCCGACAGCAGAATG-3’<br>Reverse: 5’-GCAGAAGAGGTAGCCAAACACC-3’ |
| *Sdha* | Forward: 5’-GCCATCCATTACATGACAGAGC-3’<br>Reverse: 5’-CACACAGCAACACCGATGAG-3’ |
| *Ldha* | Forward: 5’-CAGGCTCCCCAGAACAAGAT-3’<br>Reverse: 5’-CAACAAGGGCAAGCTCATCC-3’ |
| *Hdac4* | Forward: 5’-CATGGGTACTGCTGTAGGGG-3’<br>Reverse: 5’-ATGAGCTCCCAAAGCCATC-3’ |
| *Hdac6* | Forward: 5’-GGAGACAACCCAGTACATGAATGAA-3’<br>Reverse: 5’-CGGAGGACAGAGCCTGTAG-3’ |
| *Hdac9* | Forward: 5’-CCAAGTCACTGGGGCATCTT-3’<br>Reverse: 5’-TGTTCCTCTCCCAGGGTTCT-3’ |
| *Hdac10* | Forward: 5’-GGCATCGCTGAATGAGTACA-3’<br>Reverse: 5‘-GGATGAGGATCTTGCCACAC-3’ |
| *Hdac11* | Forward: 5’-GGGGGATCTCAGTGATGGTA-3’<br>Reverse: 5’-AAGAGAAGCTGCTGTCCGAT-3’ |
| *huGapdh* | Forward Primer: 5’ AGTCAGCCGCATCTTCTTTT-3’<br>Reverse Primer: 5’ CCCAATACGACCAAATCCGT 3’ |
| *huIrg1* | Forward primer 5’ TTCCGTGGTAGGAACGTTGG 3’<br>Reverse primer 5’ CTCGGCACTTTGTCGAGCTA 3’ |

#### 2.2 Supplementary Figures

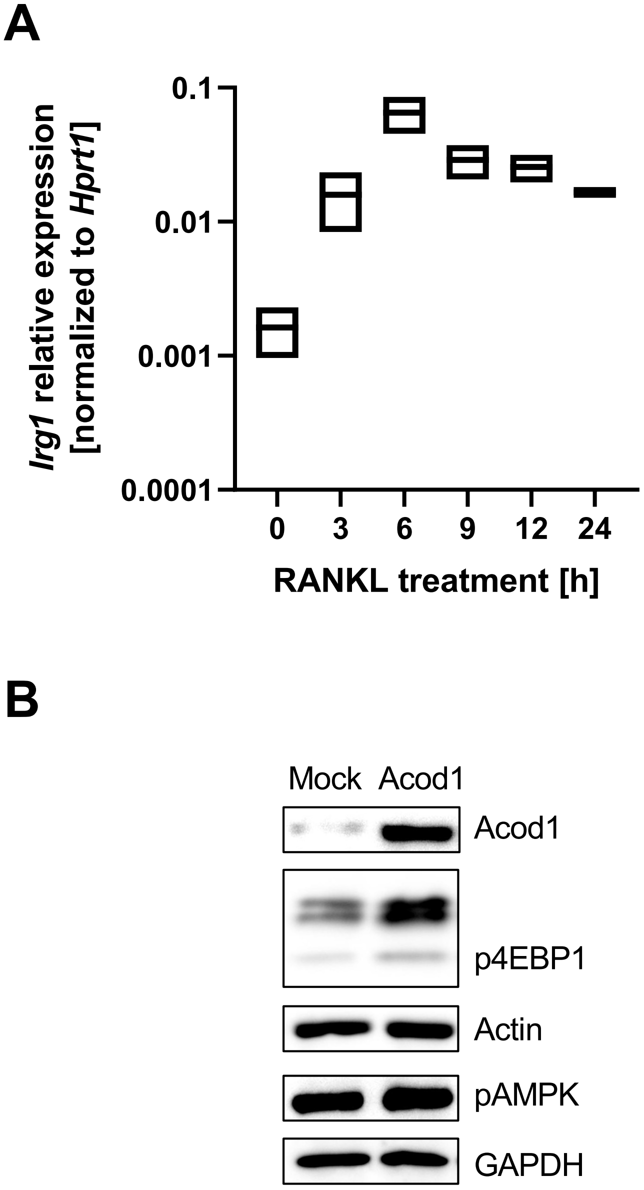


**Fig. S1 (A)** *Irg1* gene expression analysis by qPCR. RAW264.7 macrophages were stimulated with 50 ng/mL RANKL as the indicated time-points. Relative Irg1 expression levels were normalized to the reference gene *Hprt1*(n=2). **(B)** Western Blot analysis of the metabolic markers phospho-AMPK and phospho-4EBP1. Mock and ACOD1 cells were seeded and harvested on d1. Acod1 was included as a verification of the Acod1 overexpression. Actin and GAPDH are shown as the respective controls to ensure equal loading.

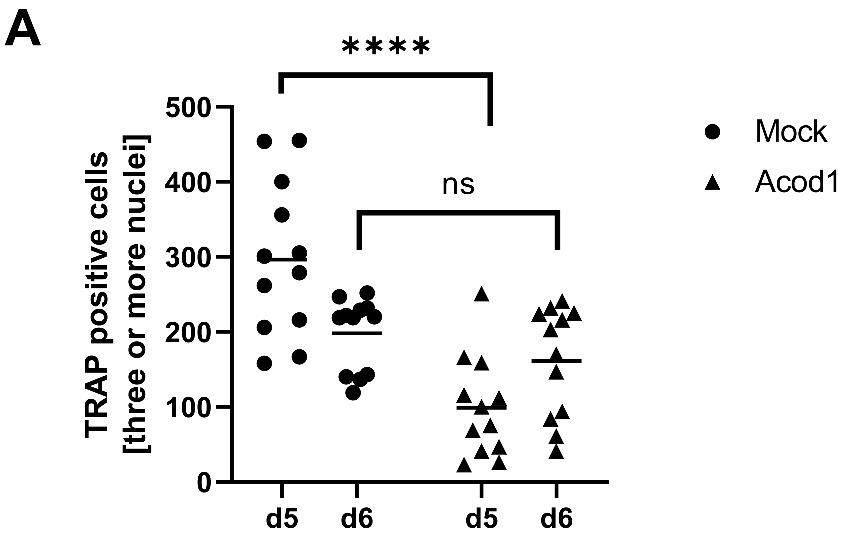


**Fig. S2** Quantification of TRAP staining. Both Mock and ACOD1 cells were seeded as duplicates and stimulated with 50 ng/mL RANKL. TRAP staining was performed after five and six days of differentiation (n=6). Statistical analysis was performed using Mann-Whitney U test.

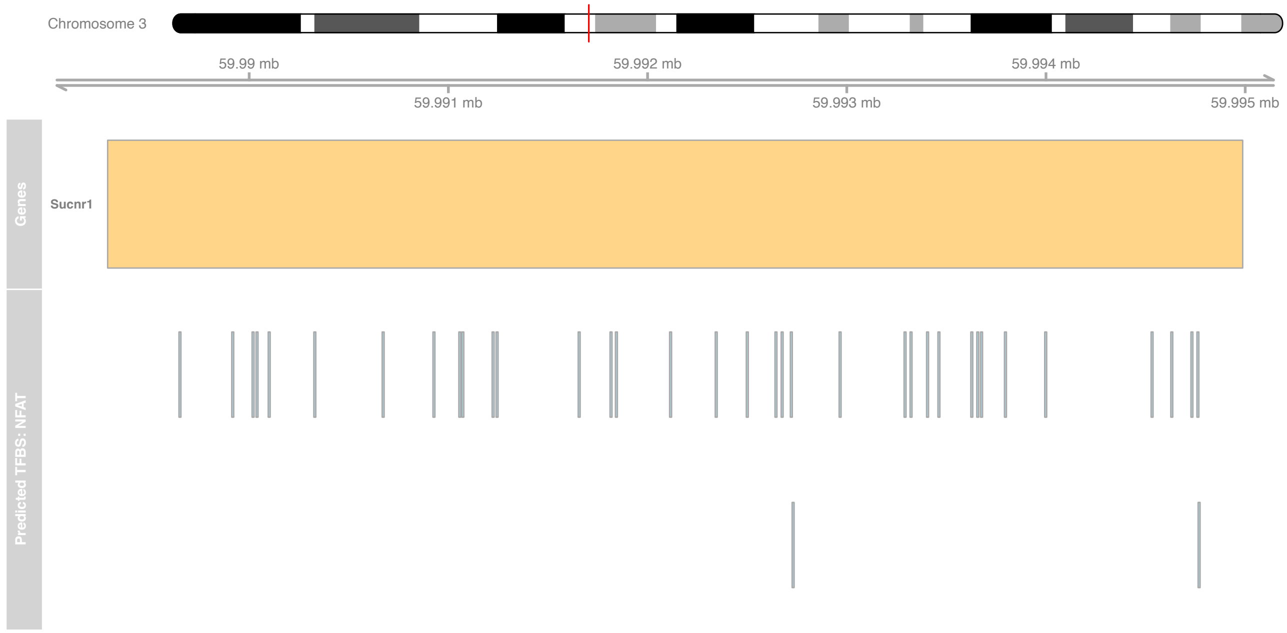


**Fig. S3** Bioinformatical analysis of the *Gpr91* promoter region for NFAT binding sites. In total, 1.608 predictions of various transcription factor binding sites were found for the *Gpr91* sequence.

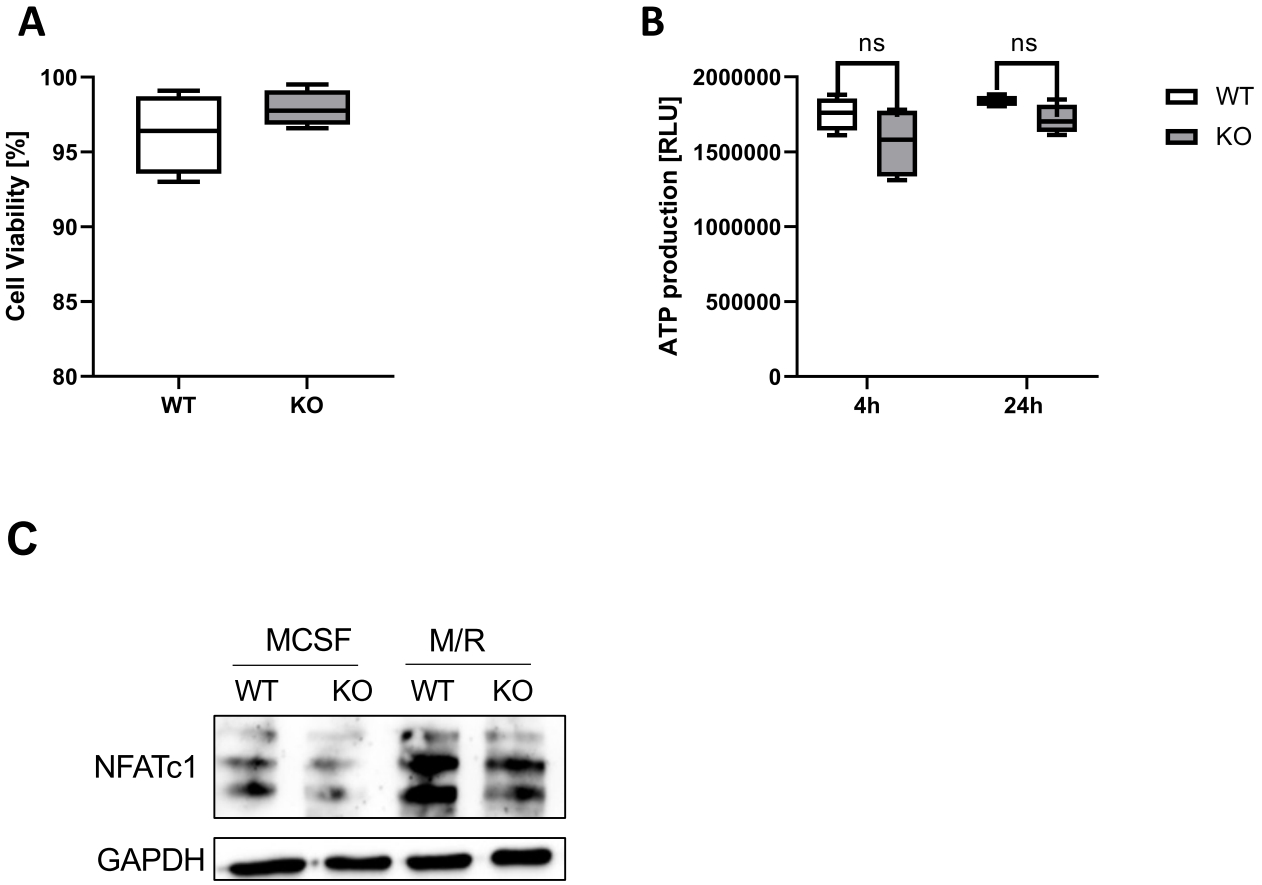


**Fig. S4 (A)** Cell viability of WT and KO cells was assessed by Zombie NIR™ staining. The percentage of live cells was shown (n=4). **(B)** The evaluation of ATP content in WT and ACOD1$^{-/-}$ BMDMs. The amount of ATP production of cells was assessed by Cell TiterGlo® Assay. Same number of WT and KO cells were seeded and evaluated after 4 or 24 h cultivation (n=4). **(C)** Western Blot analysis id NFATc1 expression in wildtype and ACOD1-deficient bone marrow cells with and without RANKL treatment

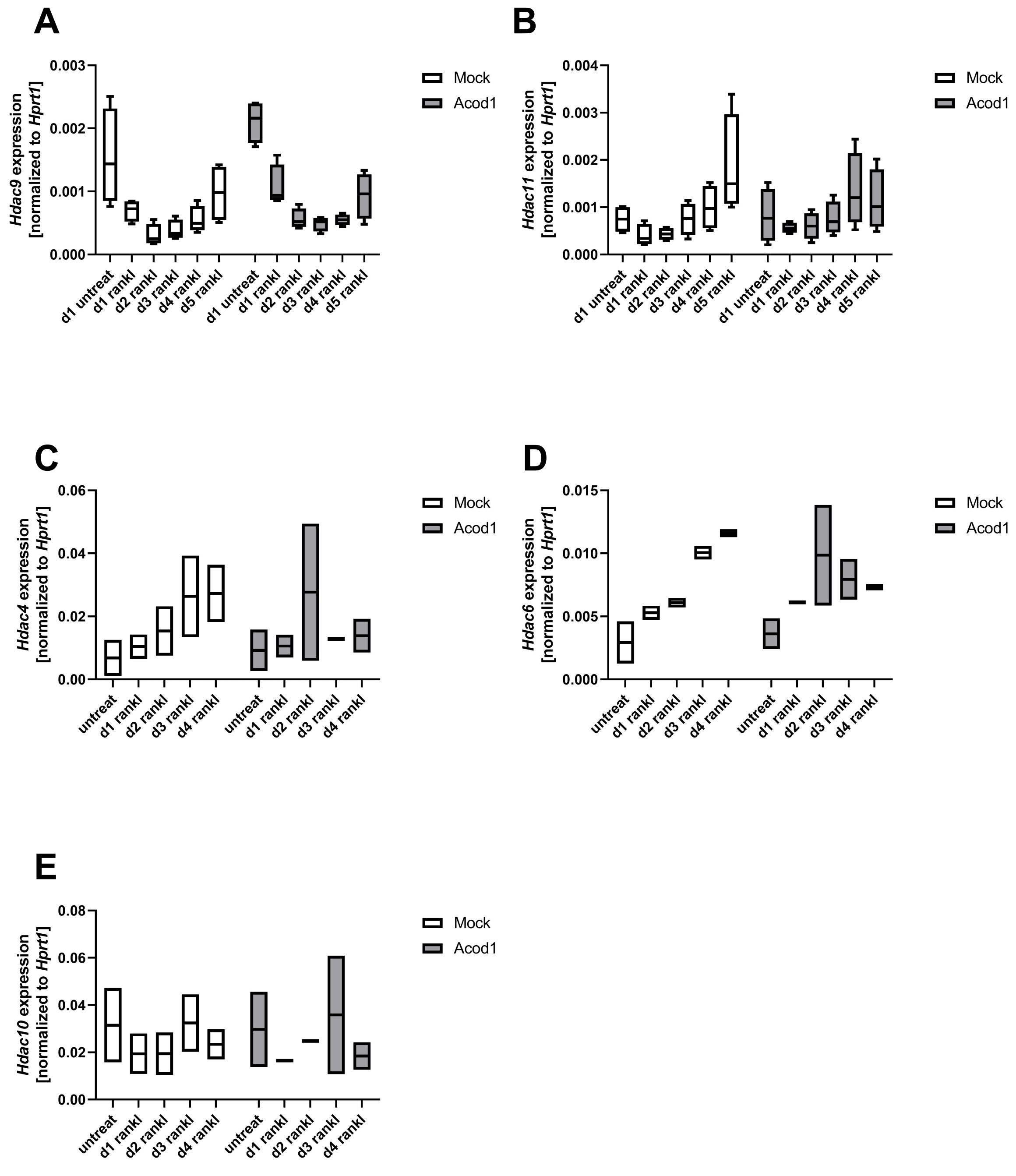


**Fig. S5** Gene expression analysis of *Hdac9* **(A)**, *Hdac11* **(B)**, *Hdac4* **(C)**, *Hdac6* **(D)**, and *Hdac10* **(E)** Cells were stimulated with 50 ng/mL RANKL and harvested as the indicates time-points. The relative gene expression was normalized to the reference gene *Hprt1*.
